# Structural Basis of Conformational Dynamics in the PROTAC-Induced Protein Degradation

**DOI:** 10.1101/2024.01.02.572291

**Authors:** Hongtao Zhao

## Abstract

Pronounced conformational dynamics is unveiled upon analyzing multiple crystal structures of the same proteins recruited to the same E3 ligases by PROTACs, and yet, is largely permissive for targeted protein degradation due to the intrinsic mobility of E3 assemblies creating a large ubiquitylation zone. Mathematical modelling of ternary dynamics on ubiquitylation probability confirms the experimental finding that ternary complex rigidification need not correlate with enhanced protein degradation. Salt bridges are found to prevail in the PROTAC-induced ternary complexes, and may contribute to a positive cooperativity and prolonged half-life. The analysis highlights the importance of presenting lysines close to the active site of the E2 enzyme while constraining ternary dynamics in PROTAC design to achieve high degradation efficiency.

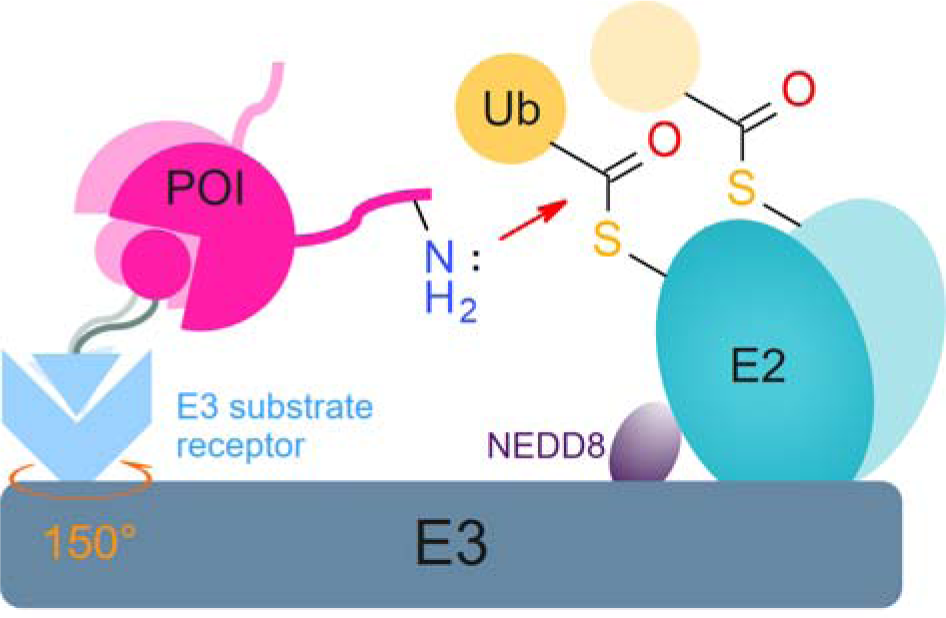

## INTRODUCTION

Targeted protein degradation (TPD) is an emerging therapeutic modality with the potential to tackle disease-causing proteins that have historically been considered undruggable due to intractable functional sites.^1^ Proteolysis-targeting chimeras (PROTACs) are heterobifunctional small molecules comprising one binder to an E3 ubiquitin ligase (E3), the other to a protein of interest (POI) and a linker connecting the two binders. Recruitment of the POI in proximity to an E3 ligase by the PROTAC triggers ubiquitylation of the POI and its subsequent degradation by the ubiquitin–proteasome system (UPS). Harnessing the advantages of catalytic degradation over inhibition^2, 3^ as well as the potential to broaden the druggable proteome,^4^ it is the most rapidly evolving field of innovation in small molecule drug discovery with more than 20 PROTACs currently in clinical trials.^5^

Ternary complex formation, which could be positively or negatively cooperative,^6, 7^ is a critical step in the mechanism of action of PROTACs by hijacking the UPS for targeted protein degradation. Negatively cooperative PROTACs can efficiently induce protein degradation too.^7^ However, not every ternary complex resulted in appreciable protein degradation, due to both kinetic and spatial aspects of the ubiquitylation process.^8–10^ Based on the SPR-measured dissociative off-rates, the lack of observed degradation was attributed to short residence time of the ternary complexes to allow for the ubiquitylation to occur.^8^ In contrast, accessibility of lysines for ubiquitin transfer was proposed to account for the distinct substrate specificity of two PROTACs upon modelling their induced ternary complexes which showed differential orientations of the POI recruited to the E3 ligase.^9^

Given the crucial role of ternary complex formation, several computational approaches have been developed based on rigid protein-protein docking coupled with extensive conformational sampling of PROTACs linkers.^11–14^ Computational modelling typically reveals a broad distribution of ternary complexes that include and extend beyond the crystallographically captured poses. The number of modelled ternary complex ensembles, as a proxy for the conformational dynamics of a given ternary system, was reported to have a positive correlation with cellular degradation.^14^ The conformational dynamics of a ternary system was indeed confirmed by hydrogen-deuterium exchange-mass spectrometry (HDX-MS)^15^ and HSQC experiments as well.^16^ The experiments further suggested that biologically relevant complexes well not exhibit the largest interaction surfaces between proteins.^15^ To discriminate between productive and unproductive ternary complexes, the whole CRL ligase complex was modelled to predict target protein ubiquitination based on the proximity of the target surface exposed lysine residues to the C-terminal of ubiquitin.^17–19^ While avidity and cooperativity were shown to drive protein degradation,^20, 21^ it was also known that increased ternary complex stability or rigidity need not correlate with increased degradation efficiency,^16^ revealing the complicated role of conformational dynamics.

In this work, we looked into crystallographically determined ternary structures deposited in the Protein Data Bank (PDB), with the aim to understand the conformational dynamics of POIs recruited to E3 ligases, the difference of PROTAC-induced protein-protein interactions from those naturally evolved, and most importantly, if there may exist privileged interactions among PROTAC-induced ternary complexes. We further built a mathematical model to correlate the conformational dynamics of a ternary system with the ubiquitylation probability. Taken together, our work offers insights into the dynamic nature of proximity-induced ubiquitylation and, can be used to select a productive ternary complex ensemble to guide the design of novel PROTAC linkers.

## MATHMATICAL MODELLING OF UBIQUITYLATION PROBABILITY

PROTACs bring a POI lysine in proximity to the active site of the ubiquitin-loaded E2 enzyme assembled onto the E3 ligase for ubiquitin transfer (**Figure 1A**). The ubiquitylation zone was estimated to have a dimension of up to 340 Å × 110 Å × 30 Å upon rotation of the ligase arm of CUL4 around DDB1 and CRBN with the center of rotation near the thalidomide-binding site.^22^ The analysis of 11 DDB1 crystal structures together with solvation energetics of rotation established a ubiquitylation zone restricted to 60-80 Å.^23^ The cullin-RING ubiquitin ligases (CRLs) by default are inactive, and become activated when NEDD8 is covalently attached to the C-terminal winged-helix-B (WHB) domain.^24–26^ The attachment of NEDD8 shifts the spatial distribution of the E2 enzyme toward the substrate, thereby enhancing ubiquitin transfer.^27^

**Figure 1.**
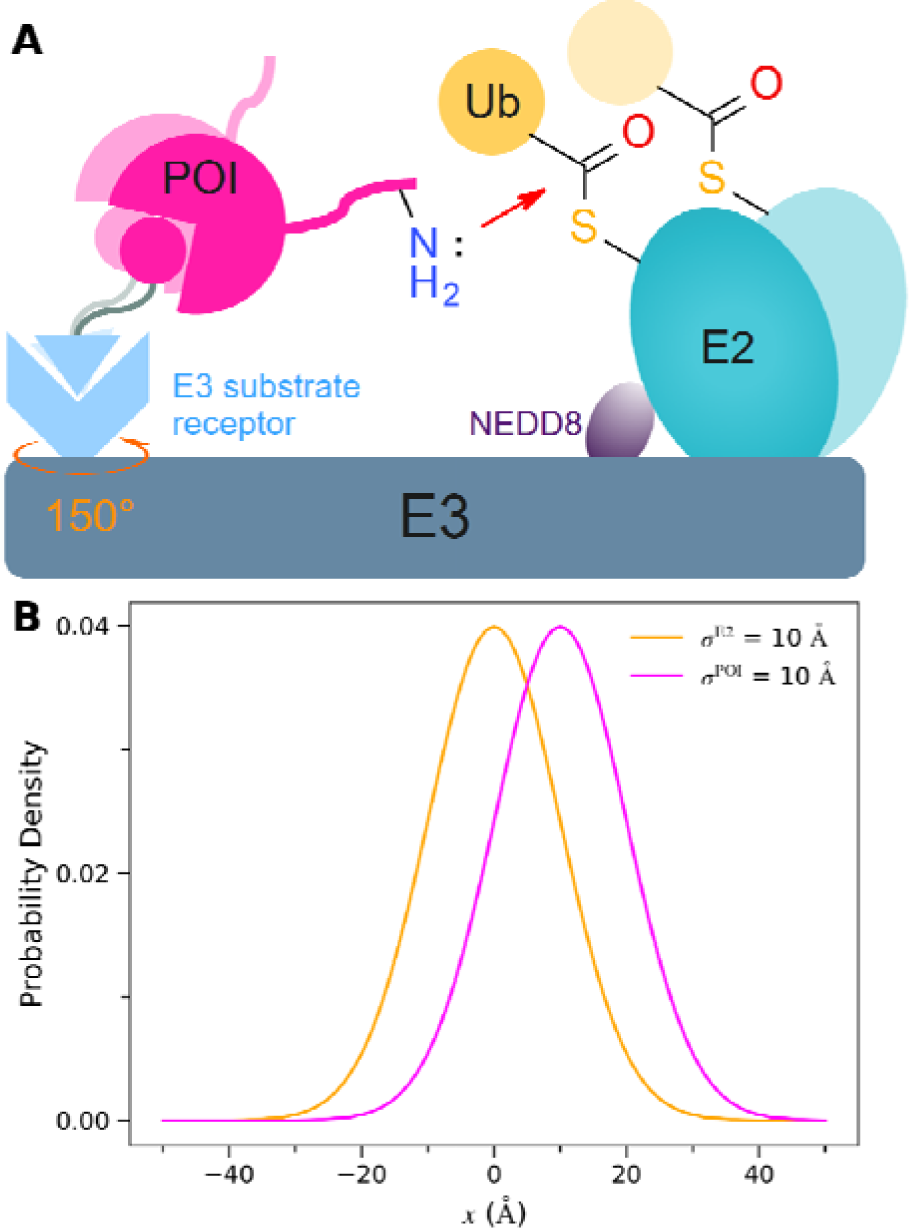
The dynamic nature of PROTAC-induced ubiquitylation. (**A**) Schematic view of the PROTAC-induced ubiquitin transfer. (**B**) Illustrative distribution of the E2 active site as well as of the POI lysines. The E3 ligase arm could rotate by at least 150° along the axis defined by the E3 substrate receptor, creating a large ubiquitylation zone.

Ubiquitylation often occur in the unstructured regions of the POI that extend beyond the globular domain anchored to the E3 ligase.^22, 23^ The surface lysines in the unstructured regions are flexible in nature. This would allow variable positioning of attacking lysines, and might as well allow a growing substrate to loop out of the active site.^28^ Furthermore, their mobility, relative to the E3 substrate receptor, could be greatly amplified by perturbations in the PROTAC binding mode, since the POI is pivoted onto the E3 substrate receptor via the PROTAC. The spatial distributions of both the E2 active site and the POI lysines follow the Boltzmann distributions governed by their underlying free energy profiles. Ubiquitylation is determined by the probability of a POI lysine found within a certain distance of the E2 active site so that nucleophilic attack of the E2-ubiquitin thioester bond by the ε-amino group of a lysine could take place.

In an effort to understand how ubiquitylation is impacted by distribution of the POI lysines as a result of the conformational dynamics of the ternary complex, we reduce the spatial distributions to one-dimensional normal distributions *N*(μ, σ) which are supported by the pair-distribution function derived from the small-angle X-ray scattering (SAXS) analysis (**Figure 1B**).^24^ The mean position of the E2 active site, relative to the E3 substrate receptor, is taken as the origin. The variance, as a measure of the spatial mobility, is fixed at 100 Å^2^ (i.e., σ^E2^ = 10 Å). The probability (*P*(*d* ≤ θ) of finding a lysine ε-amino group within a distance (θ) of the E2 active site is given by

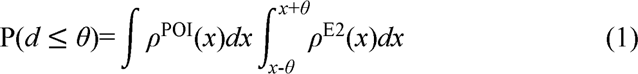

wherein *d* is the shortest distance between the POI lysine ε-amino groups and the E2 active site accounting for multi-site polyubiquitylation,^23^ ρ^POI^ and ρ^E2^ are the distribution of the POI lysine ε-amino groups and the E2 active site, respectively. The distance cutoff θ is set to 2 Å, approximating that at the transition state of the nucleophilic reaction.

Recently, we derived physical formulas of the half-maximal degradation concentration (DC_50_) from kinetic modelling of PROTAC-induced protein degradation.^29^

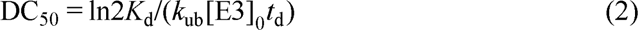

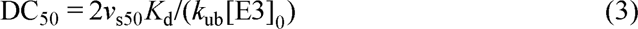

wherein *K*_d_ is the ternary affinity, [E3]_0_ is the concentration of the E3 ligase, *t*_d_ is the time of treating cells with the PROTAC before degradation of the POI was determined, *v*_s50_ is the recovery rate of the target protein at the steady state at the half-maximal degradation, and *k*_ub_ is the single effective ubiquitylation rate constant by focusing on E2-E3 ubiquitylation of the substrate where polyubiquitylation is processive and fast relative to the first monoubiquitylation.^30, 31^ The *k*_ub_ is a function of the ubiquitylation probability defined in **Eq. 1**. The **Eq. 2** describes DC_50_ measured in the initial degradation phase, and **Eq. 3** corresponds to that determined at the steady state.

## EXPERIMENTAL SECTION

### Curation of the Ternary Complex Structures

The 18 ternary complex structures collected in the PROTAC-DB 2.0 (2023)^32^ were complemented by 15 additional structures available in PDB, yielding a total of 33 complex structures for analysis. The full list is in **Table S1**.

### Buried Surface Area

The surface area buried at a protein-protein interface was calculated as the sum of the solvent accessible surface area of each monomer (i.e., the POI and the E3 ligase) minus the solvent accessible surface area of the complex in the absence of the PROTAC. The calculations were performed by the program FreeSASA^33^ using the Lee-Richard algorithm^34^ with 300 slices per atom and water represented as a sphere of radius of 1.4 Å.

### Hydrophobicity Calculation

The buried surface area of each interface was further analysed to determine the hydrophobic content by treating all carbons as apolar and the other atoms as polar. The interface hydrophobicity was then calculated as the percentage of buried surface area composed of carbon atoms only.

## RESULTS AND DISCUSSION

### Conformational Dynamics of POIs Recruited to E3 Ligases

The 33 ternary complexes from PDB (accessed on 2023.09) correspond to three E3 ligases, namely Von Hippel-Lindau (VHL), Cereblon (CRBN) and Baculoviral IAP repeat-containing protein 2 (BIRC2), each having 25, 5 and 3 structures, respectively (**Figure 2**). Despite the nearly 600 E3 ligases predicted in the human genome,^35, 36^ these three E3 ligases have been most widely co-opted to date, and have provided critical insights into the nature of PROTAC-mediated target recruitment. Notably, PROTACs in clinical phases hinge on recruiting either VHL or CRBN.^5, 37^ The two bromodomains in BRD4 were most crystalized having 13 ternary complexes (8 structures paired with VHL and 5 with CRBN),^6, 7, 38–41^ followed by the bromodomain of SMARCA2 (7 with VHL),^18, 21, 42, 43^ WDR5 (6 with VHL),^44, 45^ BTK (3 with BIRC2),^16, 46^ the bromodomain of SMARCA4 (2 with VHL),^21, 42^ FAK (1 with VHL),^47^ and BCL-xL (1 with VHL).^48^

**Figure 2.**
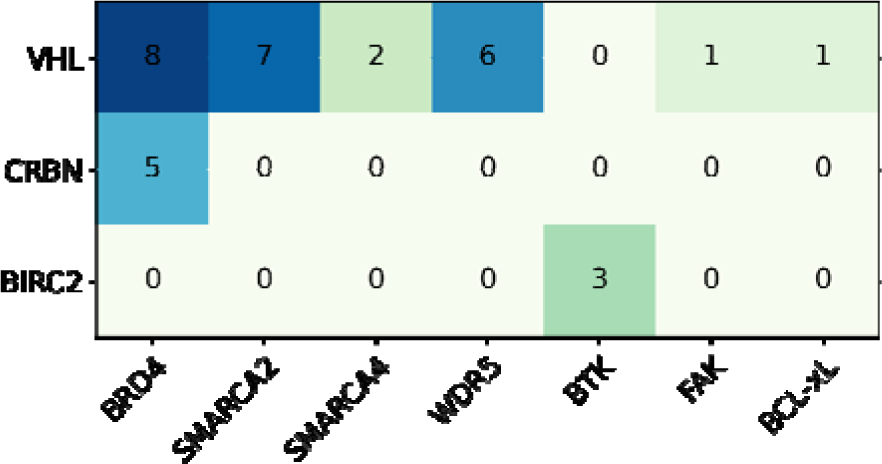
Heatmap representation of the analyzed crystal ternary complexes.

The multiple structures of the same target protein recruited to one identical E3 ligase by a variety of PROTACs help understand the conformational dynamics of a ternary system. Pronounced conformational dynamics are observed on SMARCA2, SMARCA4 and WDR5 recruited to VHL, as well as BTK to BIRC2 (**Figure 3**). Either of the two bromodomains of BRD4, termed BD1 and BD2, could be recruited to CRBN or VHL by a PROTAC. Among the five CRBN-BRD4^BD1^ ternary complexes, four structures (PDB entry 6boy, 6bn7, 6bn8 and 6bn9) showed nearly identical orientations of the BRD4^BD1^ recruited to CRBN, despite the four PROTACs having different linkers or exit vectors (**Figure S1**). In contrast, one structure (6bnb) showed a remarkable reorganization of the C-terminal domain (CTD) of CRBN relative to the N-terminal domain (NTD) and the helical-bundle domain (HBD), suggesting the large intrinsic conformational plasticity of CRBN. Concomitantly, the BRD4^BD1^ was recruited to the CRBN-CTD in an orientation distinct from that shown in the other four structures. Both BD1 and BD2 of BRD4 could be recruited to VHL in a nearly identical orientation, respectively, by a set of PROTACs (**Figure S2**). Strikingly, the binding orientation of BD1 and BD2 is completely opposite to each other. Collectively, there exists large conformational dynamics of POIs recruited to E3 ligases.

**Figure 3.**
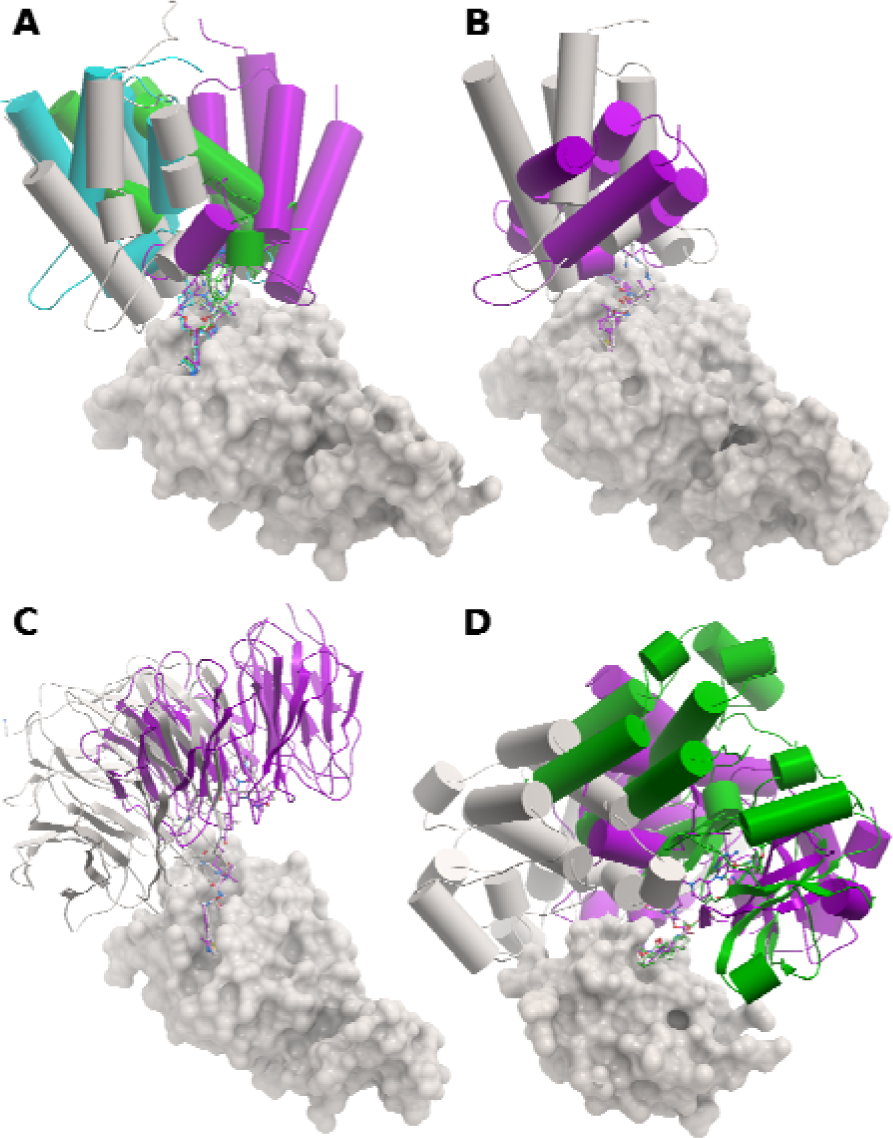
Conformational dynamics revealed by distinct orientations of POIs recruited to E3 ligases. (**A**) SMARCA2 bound to VHL (PDB entry 8g1p in grey, 7z77 in cyan, 7z6l in green and 7z76 in magenta). (**B**) SMARCA4 bound to VHL (6hr2 in grey and 8g1q in magenta). (**C**) WDR5 bound to VHL (7jtp colored in grey and 7q2j in magenta). (**D**) BTK bound to BIRC2 (6w7o in grey, 6w8i in green and 8dso in magenta). Structures were aligned on their respective E3 ligases using backbone atoms. E3 ligases and POIs were shown in the surface and cartoon representations, respectively.

Formation of a ternary complex is driven by the high affinity of a PROTAC toward both the POI and E3 ligase. Consequently, the POI could adapt to a wide range of surface patches around the binding site of the E3 ligase, even at the cost of negative cooperativity.^7^ The PROTAC-mediated protein-protein interactions are thus distinct from naturally evolved or molecular glue modulated protein-protein interfaces. Intriguingly, ternary conformational dynamics or binding plasticity is largely permissive for targeted protein degradation, given that only one analysed ternary complex resulted in no appreciable protein degradation (**Table S1**). The future ubiquitinomics analysis could help map out spatial hot-spots of lysines accessible for ubiquitin transfer in the ubiquitylation zone of an E3 ligase assembly. Together with measured ternary affinities, the analysis by **Eq. 2**/**3** could further deconvolute which ternary conformation is most effective for ubiquitylation.

### Modelling of Conformational Dynamics on the Ubiquitylation Probability

Given the experimentally observed large conformational dynamics of POIs binding to E3 ligases, we set out to have a mathematical understanding of its impact on the ubiquitylation probability. The mean shortest distance between any POI lysine and the catalytic site of the E2 enzyme (μ^POI-E2^) was simulated in the range between 2 and 25 Å, with a standard deviation (σ^POI^) up to 40 Å (**Figure 4**). The simulation results suggest that conformational dynamics is largely permissive for ubiquitylation with the probability ranging from 1% to 16%, in agreement with the crystallographically observed plasticity of POIs binding to E3 ligases. The conformational dynamics of a ternary system is permitted for ubiquitylation by the intrinsic mobility of an E3 assembly, which was revealed by the SAXS analysis^24^ and cryo-EM densities^27^, creating a large ubiquitylation zone.^22, 23^

**Figure 4.**
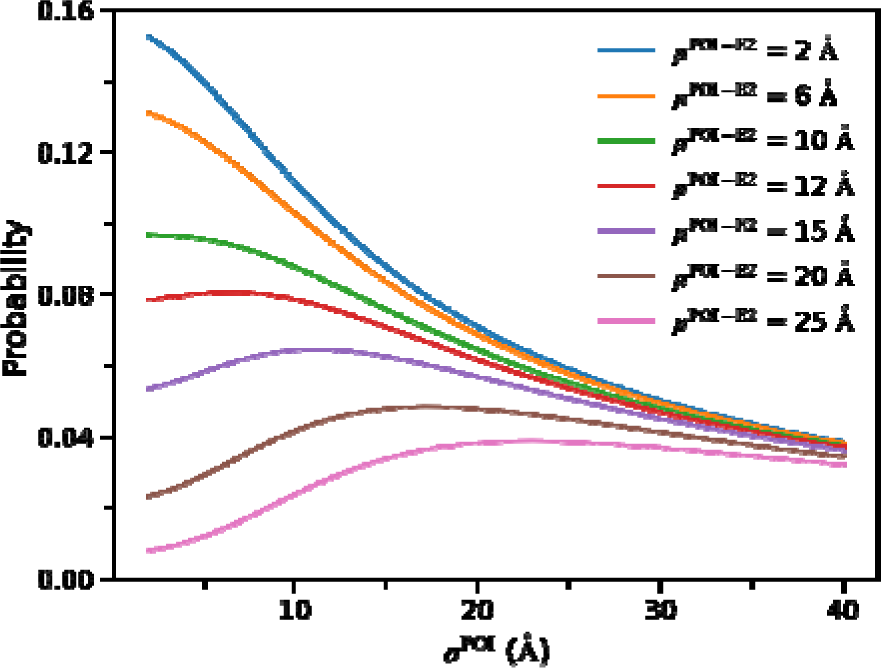
Modelling of conformational dynamics on the ubiquitylation probability.

Briefly, the increase in conformational dynamics of a ternary system leads to a monotonic decrease in the ubiquitylation probability when the mean shortest distance of μ^POI-E2^ is within 10 Å, suggesting that rigidification of the ternary complex would be beneficial for induced protein degradation. When μ^POI-E2^ is greater than 10 Å, counterintuitively, the ubiquitylation probability firstly rises to a peak, and then declines with the increase in conformational dynamics, suggesting that increased ternary complex rigidity need not always correlate with increased degradation efficiency.

### PROTAC-Induced Protein-Protein Interactions

Next, we sought to understand the difference of PROTAC-induced protein-protein interactions from those naturally evolved. The calculated buried surface areas between POIs and E3 ligases were centered around 700 Å^2^ ranging from 100 Å^2^ to 1100 Å^2^ (**Figure 5**). The buried surface areas from a nonredundant set of 144 evolved protein-protein complexes with *K*_d_ ranging between 10^−5^ and 10^−14^ M, on the other hand, has a mean value of 1817 Å^2^ ranging from 808 Å^2^ to 6254 Å^2^.^49^ This comparison indicates that PROTAC-induced ternary complex formation is largely driven by high affinities of both warheads toward their respective proteins, rather than by protein-protein interactions. Major protein conformation changes are present in most of the evolved complexes, and large movements or disorder-to-order transitions are frequently observed.^49^ In contrast, the formation of PROTAC-induced complex is characterized by rigid-body associations except for one case where the C-terminal domain of CRBN was rotated and shifted relative to the other two domains (**Figure S1**). This rigid-body association is consistent with much less involvement of protein-protein interactions in PROTAC-induced complexes.

**Figure 5.**
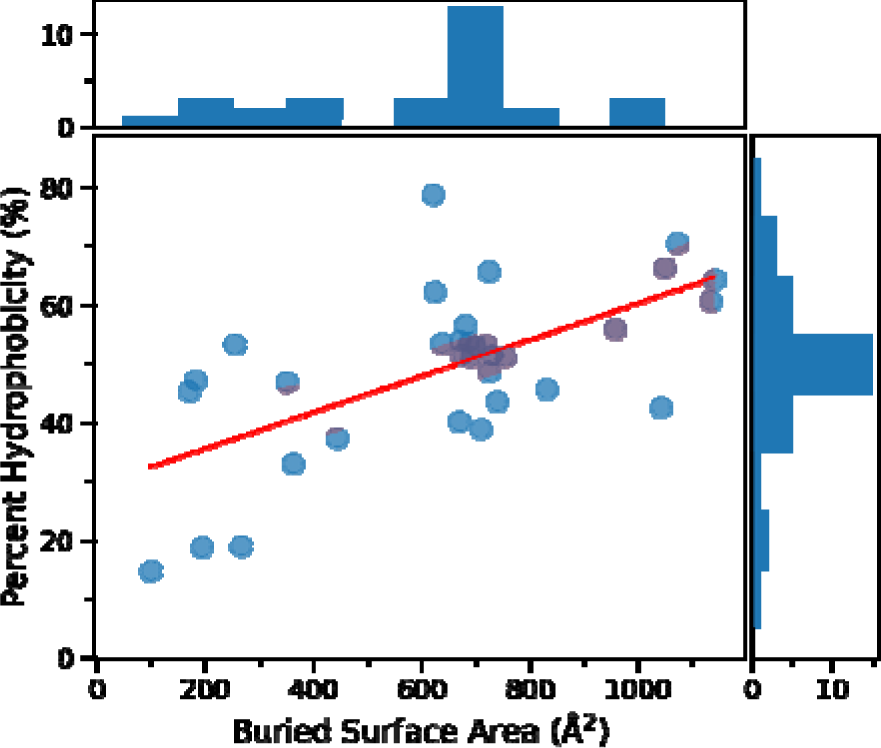
Scattering plot showing the relationship between the percent buried hydrophobicity and the buried surface area of PROTAC-induced protein-protein interactions. Their respective distributions were shown on the top and right.

The percent hydrophobicity of buried surface areas was peaked at 50% ranging from 10% to 80% (**Figure 5**). There appears to be a weak trend toward lower percent hydrophobicity for smaller buried surface areas. That means small buried surface areas might be characterized by polar interactions, which contribute to slow off-rates by functioning as kinetic traps.^50^ Given the fundamental role of the ternary complex lifetime,^8^ we focused on salt bridges which are formed by spatially proximal pairs of oppositely charged residues between POIs and E3 ligases. Salt bridges are the strongest molecular interactions and known to constrain flexibility and motion.^51^ Remarkably, 68% of the PROTAC-induced ternary complexes have at least one salt bridge (**Table S1**). Despite sharing 50% sequence identity, the two bromodomains of BRD4, BD1 and BD2, bind to VHL in completely opposite orientations, which are dictated by salt bridges (**Figure 6**). Superposition of the BRD4^BD2^ on the BRD4^BD^^1^ suggests that BD2 could establish similar interactions with VHL as BD1 including the salt bridge between its Glu438 with Arg107 from VHL. However, in the native mode of BD2 bound to VHL, in addition to the salt bridge between Glu438 and Arg69^VHL^, there exists an electrostatic zipper structure formed between Asp381-Arg108-Glu383-Arg107.^6^ In contrast, this electrostatic zipper structure would be abolished by the electrostatic repulsion between Lys91^BD1^ (corresponding to Ala384 in BD2) and Arg108^VHL^.

**Figure 6.**
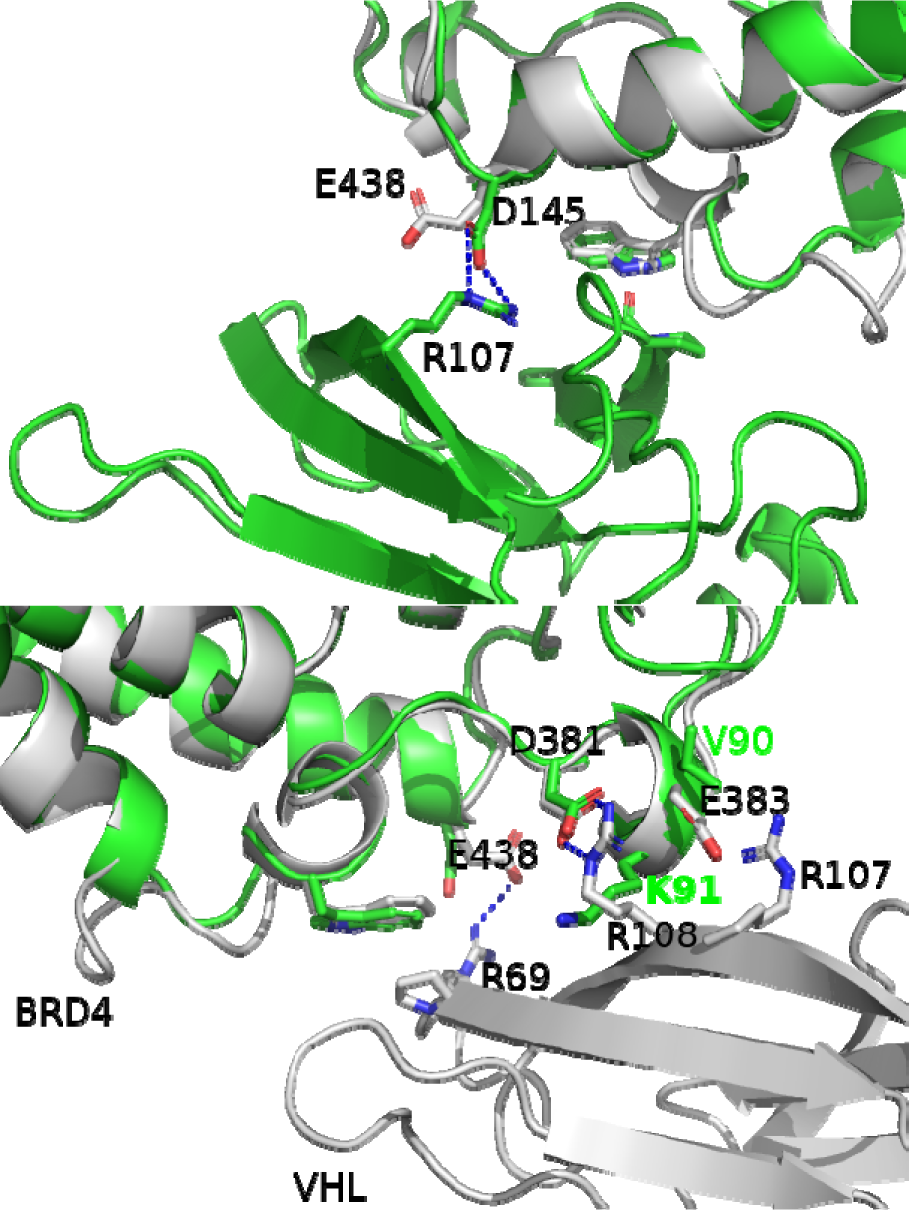
Salt bridges dictating the opposite orientations of BRD4^BD^^1^ and BRD4^BD2^ recruited to VHL. (Top) Superposition of the BRD4^BD2^ colored in grey from the PDB structure 5t35 on the BRD4^BD^^1^ of 7khh in green. (Bottom) Superposition of the BRD4^BD^^1^ of 7khh in green on the BRD4^BD2^ of 5t35 in grey.

### Implication of Conformational Dynamics in PROTAC Design

To put conformational dynamics into perspective, we looked at two pairs of PROTACs, for each of which the crystal ternary complex structure was determined (**Figure 7**). PROTAC **2** induces cellular degradation of WDR5 with a DC_50_ value of 0.05 _μ_M while PROTAC **1** shows no appreciable degradation at concentrations up to 10 _μ_M.^52^ Both compounds have rather similar inhibitory potency against WDR5 in cells or lysates, indicating that permeability is not the determining factor for the difference in degradation. Both compounds recruited WDR5 to VHL in a similar orientation. In comparison with the four-repeated PEG linker of **1**, the four-carbon alkyl linker of **2** permits WDR5 a bit close to VHL (**Figure 8**). The nuance in the binding conformations enables one salt bridge formed between Lys259^WDR5^ and Asp92^VHL^, in addition to the two H-bonds formed between Arg69 from VHL and the two backbone oxygens from WDR5. These electrostatic interactions may lead to a positive cooperativity, which contributes to a lower DC_50_ value evident from **Eq. 2/3**. The electrostatic interactions may also lead to a prolonged lifetime of the complex, which was shown to influence degradation.^8^ The effect of lifetime on the downstream reaction was recently confirmed by our Monte Carlo simulations.^53^ When the lifetime is insufficiently long to permit the downstream reaction (e.g., ubiquitylation), the downstream rate shows a linear dependency on the dissociation barrier of the complex before it reaches a plateau. The plateau means that extremely stable ternary complex is unnecessary and even hampers the catalytic turnover frequency of the PROTAC because it acts via transient binding events. The three-PEG linker was shown to be very flexible in molecular dynamics simulations of MZ1,^54^ and the four-PEG linker of **1** together with the lack of favorable protein-protein interactions suggests that the ternary structure of WDR5-**1**-VHL is rather dynamic in solution, which could be detrimental to the ubiquitylation probability (**Figure 4**). The WDR5 PROTAC pair demonstrates the beneficial effect of linker rigidification on targeted protein degradation via positive cooperativity, increased lifetime and decreased conformational dynamics.

**Figure 7.**
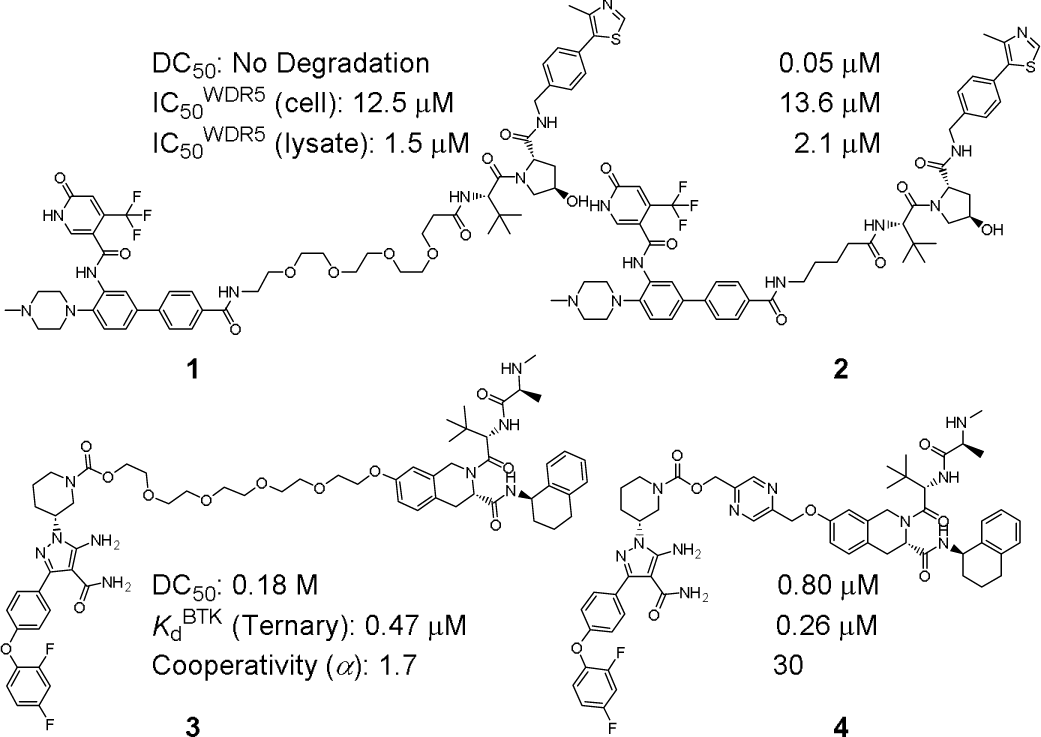
PROTACs illustrating the role of conformational dynamics in DC_50_.

**Figure 8.**
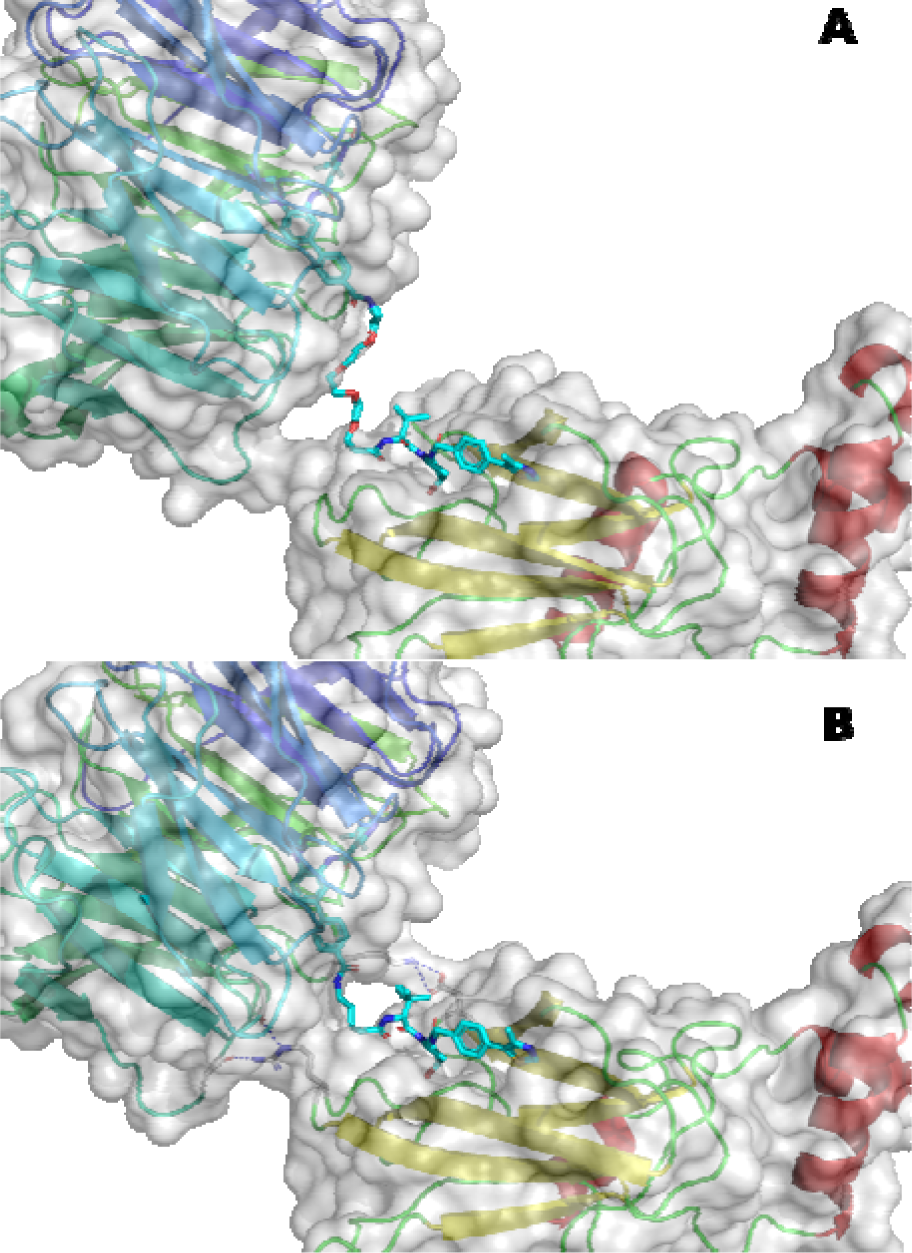
Ternary complex structure of WDR5-**1**-VHL (**A**) and WDR5-**2**-VHL (**B**). PDB entry 8bb2 and 7q2j.

Structure-enabled design has led to replacement of the central four-PEG linker of PROTAC **3** by a pyrazine, yielding a new degrader **4**.^16^ PROTAC **4** showed a 4-fold slower off rate than **3**, and a nearly 20-fold increase in cooperativity which led to a 2-fold increase in the ternary affinity against BTK. Surprisingly, it is 4-fold less potent than **3** to degrade BTK. Both compounds were calculated to have similar cellular permeability. In addition, a similar loss in ubiquitylation activity in a cell-free in vitro system was observed. The HSQC spectrum suggested a rather flexible ternary assembly of BTK-**3**-BIRC2, and three distinct ternary conformations were captured crystallographically in the asymmetric unit (**Figure 9**). In contrast, the BTK-**4**-BIRC2 complex was characterized by an overall increase in rigidity, and a reduced conformational ensemble from modelling dominated by a single, highly populated cluster. Indeed, the crystal structure of BTK-**4**-BIRC2 revealed only one ternary conformation. The modest degradation potency of both compounds suggest that the accessible lysines of BTK might be positioned, on average, away from the E2 active site. If so, the unexpected loss of degradation potency upon increasing ternary complex stability could be accounted for by our mathematical modelling shown in **Figure 4**.

**Figure 9.**
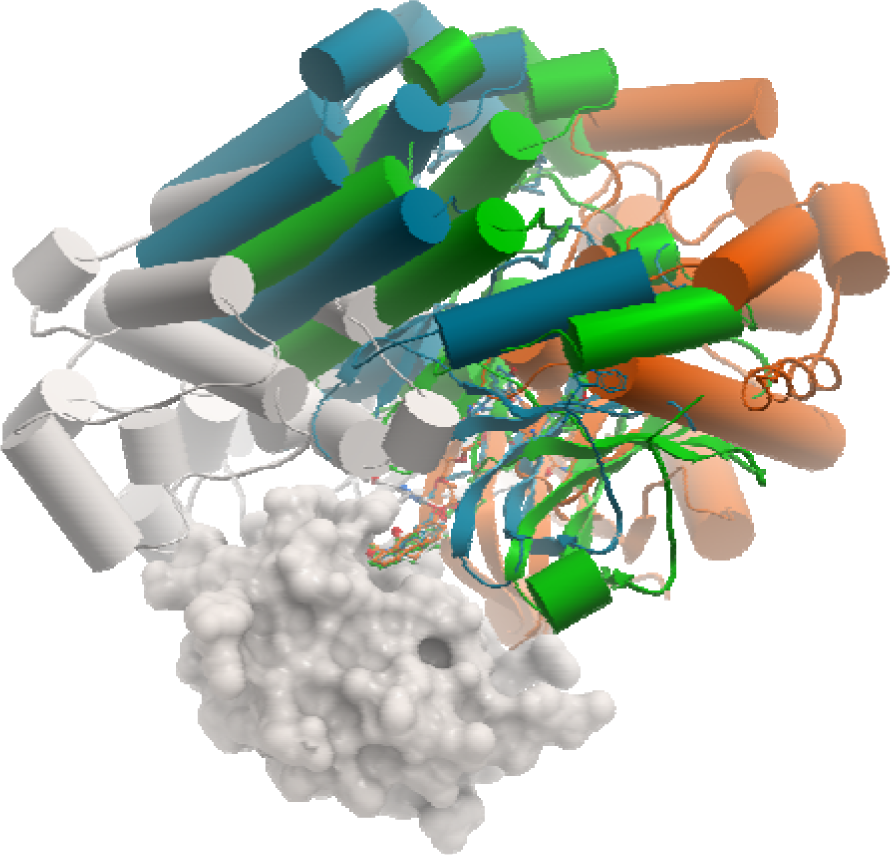
Ternary complex structure of BTK-**4**-BIRC2 (6w7o, colored in grey) superimposed on BIRC2 of the structure of BTK-**3**-BIRC2 (6w8i, the three copies of BTK in the asymmetric unit were colored in blue, green and orange, respectively). BIRC2 was shown in surface.

From the modelling perspective, it is clear that high ubiquitylation probability is only achievable when accessible lysines of POIs are positioned close to the E2 active site (**Figure 4**). This notion is in line with the recent ground-breaking computational work of predicting the proximity of POI lysines to the E2 enzyme in the context of the full cullin-RING ligase.^17, 18^ When the average distance between lysines and the E2 active site is short enough, it is beneficial to reduce conformational dynamics of the ternary system by establishing protein-protein contacts or linker rigidification.

## CONCLUSIONS

In comparison with evolved protein complexes, PROTAC-induced ternary complexes are characterized by rigid-body associations having protein-protein interactions to a lesser extent. Pronounced conformational dynamics of POIs recruited to E3 ligases are observed in the crystal ternary structures and, are largely permissive for protein degradation owing to the intrinsic mobility of E3 ligase assemblies. Mathematical modelling of ternary dynamics on the ubiquitylation probability confirms the experimental finding that ternary complex stability need not always correlate with increased protein degradation. Nevertheless, high ubiquitylation probability can only be achieved upon positioning accessible POI lysines close enough to the E2 active site, and in that case, the decrease in conformational dynamics is beneficial for protein degradation. Hence, a minimum level of protein-protein interactions could be necessary to restrain conformational dynamics of a ternary system toward protein degradation, although the formation of a ternary complex is mainly driven by high affinities of the two PROTAC warheads. Salt bridges were found to prevail in the PROTAC-induced ternary complexes, contributing to both affinity and stability.

## ABBREVIATIONS

PROTACs: proteolysis-targeting chimeras
POI: protein of interest
VHL: von Hippel-Lindau
CRBN: Cereblon
BIRC2: Baculoviral IAP repeat-containing protein 2
DC_50_: half-maximal degradation concentration.

## Notes

H. Z. is an employee of AstraZeneca and may own stock or stock options.

## Supporting Information

List of the 33 PDB entries with DC_50_ values of the corresponding PROTACs, buried surface areas, and numbers of salt bridges; conformational dynamics of BRD4 recruited to CRBN and VHL, respectively.

