## Supporting Information for "Structural Basis of Conformational Dynamics in the PROTAC-Induced Protein Degradation"

Hongtao Zhao*

Medicinal Chemistry, Research and Early Development, Respiratory and Immunology (R&I), BioPharmaceuticals R&D, AstraZeneca Gothenburg, Sweden

*

**Table S1. Statistics of the 33 ternary complex structures.**

| PDB entry | POI | E3 ligase | $DC_{50}$ (nM) | BSA (Å$^2$) | %apolar | Salt bridge |
|---|---|---|---|---|---|---|
| 6w8i | BTK | BIRC2 | 182 | 194.3 | 18.8 | 1 |
| 6w7o | BTK | BIRC2 | 800 | 1041.9 | 42.6 | 2 |
| 8dso | BTK | BIRC2 | 57 | 637.6 | 53.5 | 1 |
| 6boy | BRD4$^{BD1}$ | CRBN | 6 | 1133.43 | 60.65 | 0 |
| 6bnb | BRD4$^{BD1}$ | CRBN | 500 | 957.22 | 55.92 | 1 |
| 6bn9 | BRD4$^{BD1}$ | CRBN | 5 | 1141.46 | 64.41 | 0 |
| 6bn8 | BRD4$^{BD1}$ | CRBN | 1800 | 1072.71 | 70.53 | 0 |
| 6bn7 | BRD4$^{BD1}$ | CRBN | 50 | 1048.92 | 66.35 | 0 |
| 6zhc | BCL-xL | VHL | 4.8 | 669.5 | 40.1 | 1 |
| 7pi4 | FAK | VHL | 4 | 739.4 | 43.5 | 2 |
| 7khh | BRD4$^{BD1}$ | VHL | 0.095 | 723.9 | 65.7 | 1 |
| 8bds | BRD4$^{BD1}$ | VHL | 1100 | 624.0 | 62.3 | 1 |
| 8beb | BRD4$^{BD1}$ | VHL | 320 | 620.3 | 78.8 | 0 |
| 6sis | BRD4$^{BD2}$ | VHL | 25-125 | 680.8 | 56.5 | 1 |
| 5t35 | BRD4$^{BD2}$ | VHL | 8 | 729.6 | 51.4 | 3 |
| 7znt | BRD4$^{BD2}$ | VHL | N.A.[a] | 753.4 | 51.2 | 2 |
| 8bdt | BRD4$^{BD2}$ | VHL | 560 | 724.8 | 48.6 | 3 |
| 8bdx | BRD4$^{BD2}$ | VHL | 1100 | 690.0 | 51.2 | 3 |
| 6hay | SMARCA2 | VHL | 300 | 686.5 | 53.7 | 0 |
| 6hax | SMARCA2 | VHL | 70 | 698.3 | 52.9 | 1 |
| 7z6l | SMARCA2 | VHL | 78 | 169.9 | 45.3 | 0 |
| 7z77 | SMARCA2 | VHL | 2 | 351.0 | 46.9 | 0 |
| 7z76 | SMARCA2 | VHL | 3 | 709.3 | 38.9 | 3 |
| 7s4e | SMARCA2 | VHL | 6 | 719.2 | 53.3 | 1 |
| 8g1p | SMARCA2 | VHL | 70/221 | 673.8 | 51.5 | 0 |
| 6hr2 | SMARCA4 | VHL | 120 | 671.1 | 53.8 | 0 |
| 8g1q | SMARCA4 | VHL | N.A.[a] | 443.0 | 37.4 | 2 |
| 7q2j | WDR5 | VHL | 53 | 362.8 | 33.0 | 1 |
| 7jtp | WDR5 | VHL | 3.7 | 831.2 | 45.7 | 2 |
| 7jto | WDR5 | VHL | 260 | 266.4 | 19.0 | 2 |
| 8bb4 | WDR5 | VHL | 116 | 254.3 | 53.3 | 1 |
| 8bb2 | WDR5 | VHL | Inactive | 99.7 | 14.8 | 0 |
| 8bb5 | WDR5 | VHL | Partial | 181.1 | 47.1 | 1 |

[a] Not available.

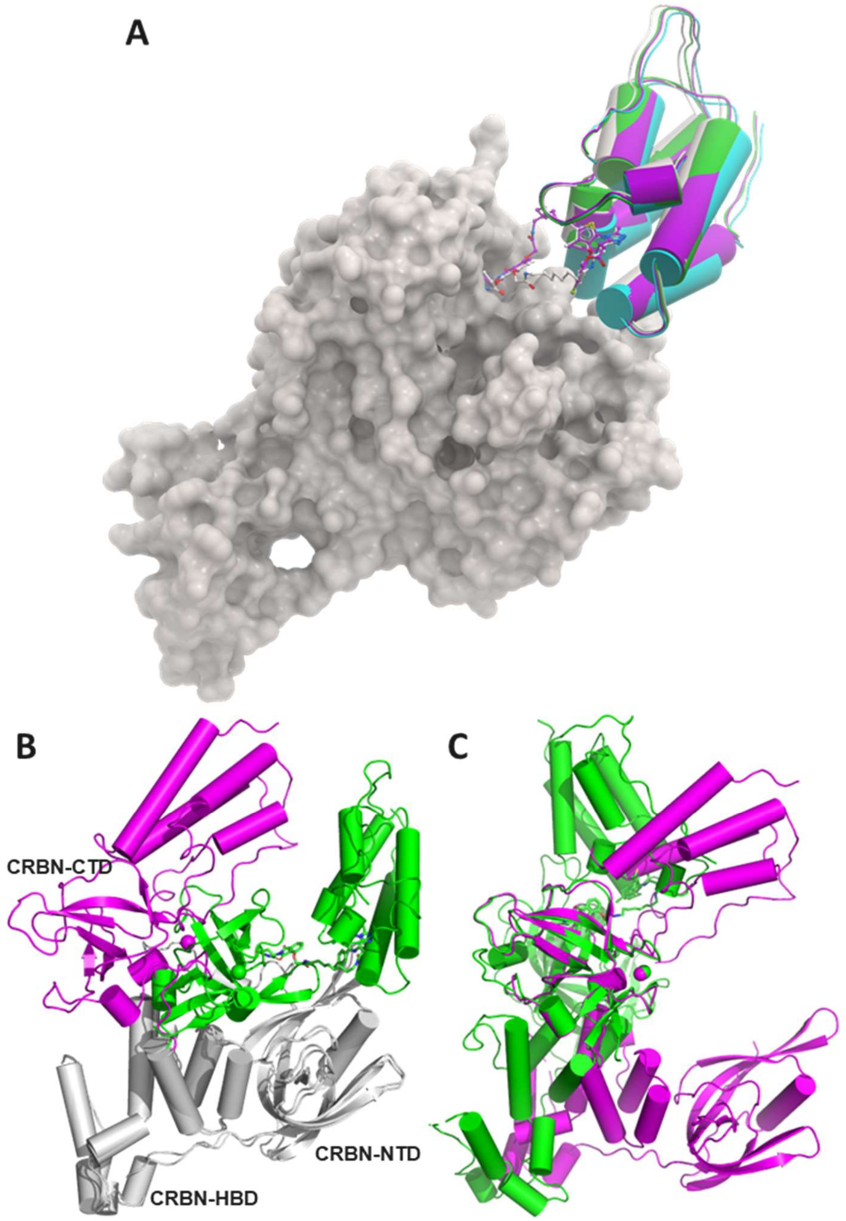


**Figure S1.** Conformational dynamics of CRBN-BRD4$^{BD1}$. (**A**) Superposition of the four CRBN-BRD4$^{BD1}$ ternary complexes (PDB entry 6boy colored in grey, 6bn9 in cyan, 6bn8 in green, and 6bn7 in magenta) over the CRBN shown as surface. (**B**) Superposition of the two CRBN-BRD4$^{BD1}$ ternary complexes (6boy with the BRD4$^{BD1}$ and CRBN-CTD in green, and 6bnb in magenta) over the CRBN-NTD/HBD colored in grey, and (**C**) over the CRBN-CTD (residues 320-400).

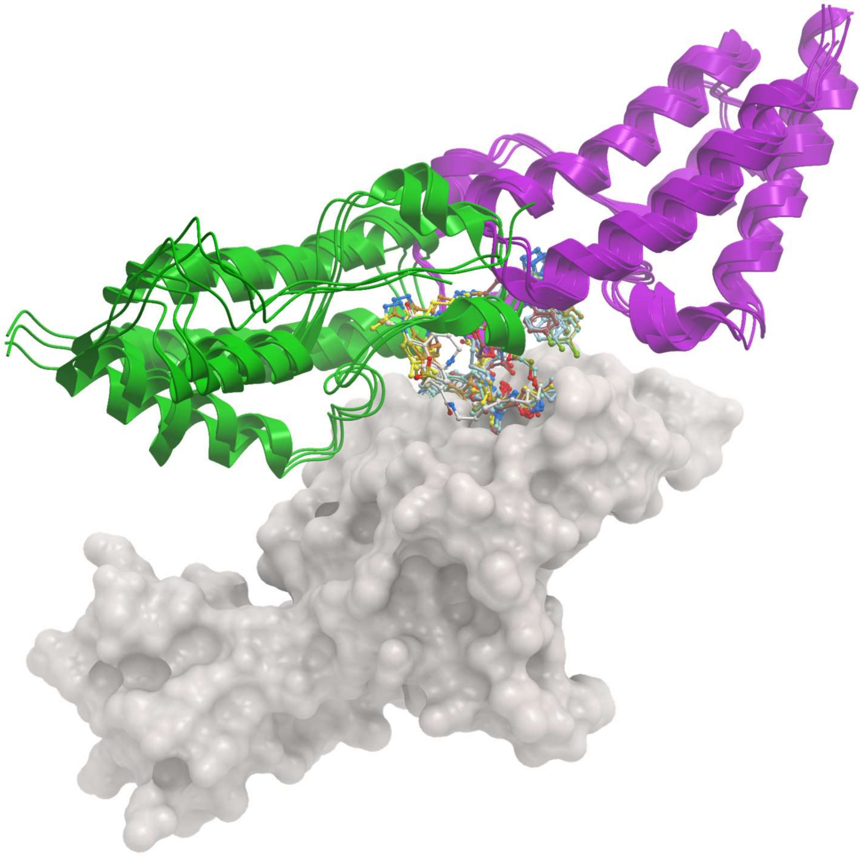

**Figure S2.** Superposition of the three VHL-BRD4$^{BD1}$ ternary complexes (PDB entry 7khh, 8bds, and 8beb) and the five VHL-BRD4$^{BD2}$ complexes (6sis, 5t35, 7znt, 8bdt and 8bdx) over the VHL shown as surface. BRD4 was shown in cartoon representation with BD1 colored in green and BD2 in magenta.
